# Compartmentalization of specialized metabolites across vegetative and reproductive tissues in two sympatric *Psychotria* (Rubiaceae)

**DOI:** 10.1101/2022.07.22.501147

**Authors:** Gerald F. Schneider, Cole A. Carlson, Elsa M. Jos, Noelle G. Beckman

## Abstract

**Premise of the study:** The specialized metabolites of plants are recognized as key chemical traits in mediating the ecology and evolution of sundry plant-biotic interactions, from pollination to seed predation. Intra- and interspecific patterns of specialized metabolite diversity have been studied extensively in leaves, but the diverse biotic interactions which contribute to specialized metabolite diversity encompass all plant organs. Focusing on two species of *Psychotria* shrubs, we investigate and compare patterns of specialized metabolite diversity in leaves and fruit with respect to each organ’s diversity of biotic interactions.

**Methods:** To evaluate associations between biotic interaction diversity and specialized metabolite diversity, we combine UPLC-MS metabolomic analysis of foliar and fruit specialized metabolites with existing surveys of leaf- and fruit-centered biotic interactions. We compare patterns of specialized metabolite richness and variance among vegetative and reproductive tissues, among plants, and between species.

**Key results:** In our study system, leaves are involved in a greater number of host-specific biotic interactions than fruit, while fruit-centric interactions are more ecologically diverse. This was reflected in specialized metabolite richness – leaves contained more than fruit, while each contained over 200 organ-specific specialized metabolites. Within each species, leaf- and fruit specialized metabolite composition varied independently of one another across plants. Specialized metabolite composition exhibited stronger contrasts between organs than between species.

**Conclusions:** As ecologically disparate plant organs with organ-specific specialized metabolite traits, leaves and fruit can each contribute to the tremendous overall diversity of plant specialized metabolites.

---

Specialized metabolites of plants constitute a major component of the ecological and evolutionary framework of plant and animal biodiversity. Through mediating plants’ ecological interactions and evolutionary relationships with consumers, parasitoids, pollinators, and seed-dispersers, specialized metabolites hold within their structural and functional diversity the potential to generate and reinforce the species richness of plants and the organisms with which they interact (Levey et al., 2007; Heil, 2008; Dicke and Baldwin, 2010; Courtois et al., 2016; Richards et al., 2015; Endara et al., 2017; Stevenson et al., 2017). To date, much of the development of theory and accumulation of evidence regarding evolutionary linkages between plant-biotic interactions and specialized metabolite diversification has stemmed from studies of plant taxa or communities and their specialist herbivores (e.g. Becerra and Venable 1999, Kursar et al. 2009, Becerra 2015, Richards et al. 2015, Endara et al. 2017, Uckele et al. 2021). However, the development of the Interaction Diversity Hypothesis (IDH) with respect to ecologically disparate plant organs (e.g. animal-dispersed fruit vs. leaves; Whitehead et al., 2021) has established the efficacy of a plant-level perspective on specialized metabolite trait evolution, one which integrates organ-level patterns and processes. In brief, the IDH posits that the diversity of specialized metabolites in a given plant species is an emergent consequence of the diversity of biotic interactions amid which the species has evolved (Berenbaum and Zangerl, 1996; Iason et al., 2011; Whitehead et al., 2021). In such a scenario, the distinct selective pressures of each antagonistic or mutualistic interaction act independently from one another on the chemical deterrents and/or attractants in the pertinent plant tissue. This scenario is hypothesized to result in a modular “tool kit” of many specialized metabolites, each suited to a distinct ecological interaction. Supporting these conceptual advances have been technological and analytical advances in the field of metabolomics, allowing quantitative and comparative evaluations of specialized metabolites at tissue-, organ-, organismic, and community scales (Wang et al., 2016; Sedio, 2017; Aron et al., 2020; Walker et al., 2022).

Overlapping with the recent refinement of the IDH, fruit and its constituent tissues have increasingly been recognized as an important component of specialized metabolite diversity, particularly in plants which rely on animals for seed dispersal (Cipollini and Levey, 1997; Whitehead et al., 2021). Fruits of zoochorous (animal-dispersed) species are subjected to exceptionally complex arrays of biotic selective pressures as compared to other organs within the same species. Specifically, zoochorous fruits are tasked with defending against numerous guilds of biotic antagonists, such as seed predators, non-dispersing pulp consumers, and fungal pathogens, while attracting seed-dispersing mutualists upon ripening (Cipollini and Levey, 1997; Whitehead et al., 2021). Considering this, the IDH supports a prediction of highly specialized metabolite diversity in fruit pericarp relative to other tissues in zoochorous plant species. This prediction has been borne out by a handful of studies, both at the scale of chemical class (Whitehead and Bowers, 2013; Whitehead et al., 2013) and that of the metabolome (Schneider et al., 2021). However, further case studies are called for (Whitehead et al., 2021) in order to establish generalities of the IDH for specialized metabolite diversity and variability across a range of biological and ecological scales.

In this study, our goals consisted of 1) broadening the taxonomic scope of IDH-based comparisons of specialized metabolite composition across vegetative and reproductive tissues, and 2) understanding variability in specialized metabolite composition across tissues and individuals with respect to species-level differences. For our focal plant taxa, we selected a pair of syntopic species of *Psychotria* (Rubiaceae), a pantropical genus of mainly shrubs and small trees in the Rubiaceae. Fruits of Neotropical *Psychotria* are bird-dispersed (Charles-Dominique, 1993; Poulin et al., 1999), and the genus likewise represents an important food source for frugivorous understory birds throughout the tropics (Snow, 1981). *Psychotria* also host a diverse assemblage of insect herbivores; for example, the 20 *Psychotria* species at Barro Colorado Island, Panama, host 127 species of insect herbivores (Sedio, 2013). With respect to specialized metabolites, *Psychotria* and closely related genera in the tribes Psychotrieae and Palicoureeae are known for their production of a structurally diverse array of alkaloids. Indeed, a chemotaxonomic approach utilizing structural groupings of Psychotrieae and Palicoureeae alkaloids has recently been employed in an attempt to disentangle the evolutionary relationships within these tribes (Berger et al., 2021).

Notably, studies of the specialized metabolites of Psychotrieae and Palicoureeae have generally been limited to vegetative tissue. However, sufficient ecological data are available to allow us to posit hypotheses regarding the specialized metabolite composition of the fruits of Psychotrieae/Palicoureeae in our focal species pair, a subset of the 20 species at Barro Colorado Island. Among these 20 species, fruits are consumed by a diverse array of birds (Poulin et al., 1999; G. F. Schneider, Utah State University, unpublished data) but are apparently unpalatable to mammalian frugivores (Wright et al., 2016). Fruit pulp and seeds of these 20 species experience low pre-dispersal pest damage (Basset et al., 2021) and are involved in a low number of antagonistic species interactions, with only 7 fruit antagonist species (Basset et al., 2021) to the 127 leaf antagonist species previously mentioned (Sedio, 2013). The avian-specific seed dispersal and low fruit damage rates are consistent with patterns observed in plant taxa known to contain distinctive specialized metabolites in fruit tissues (summarized in Levey et al., 2007), which leads us to hypothesize that *Psychotria* fruit contains distinctive specialized metabolites. Based on the IDH, the low species richness of fruit-associated antagonists (2 pulp-feeders and 5 seed predators; Basset et al., 2021) as compared to leaf-associated antagonists (127; Sedio, 2013), leads us to hypothesize that fruit pulp and seeds will exhibit lower richness of specialized metabolites, i.e. total number of distinct compounds, as compared to leaves. Finally, as specialized metabolite variability can impose behavioral and metabolic costs on consumers (Adler and Karban, 1994; Salazar et al., 2016; Massad et al., 2017; Pearse et al., 2018; Whitehead et al., 2021), we hypothesize that fruit pulp will have lower specialized metabolite variability than leaves or seeds as a result of selection to reduce costs for mutualistic consumers.

## MATERIALS AND METHODS

### Study site

Barro Colorado Island (9 9’N, 79 51’W), hereafter referred to as BCI, encompasses a lowland, moist tropical forest with average annual rainfall of 2,600 mm (Leigh et al., 1996). The rainfall on BCI arrives mainly between May and December, with a pronounced dry season between January and April (Leigh et al., 1996). Both the vegetative and reproductive phenology of the woody plants of BCI exhibit seasonality, with leaf and fruit production both peaking in the early wet season (Leigh et al., 1996) and fruit maturation peaking between the early dry season and early wet season, though the latter varies widely by species (Wright et al., 1999; Zimmerman et al., 2007).

### Study species

The two species selected for this study, *Psychotria limonensis* and *Psychotria marginata*, are distinguished from the other Psychotria of BCI–and linked to one another–by their reproductive phenology. Though only distantly related within the genus (Sedio et al., 2012), these species both undergo fruit maturation in the early wet season, while all other Psychotrieae and Palicoureeae of BCI produce fruit in the late wet season (Poulin et al., 1999). Thus, *P. limonensis* and *P. marginata* represent replicates of a distinct point of ecological intersection between *Psychotria* and leaf- and fruit-consuming invertebrates and pathogens. Both species have been included in surveys of insect herbivores (Sedio, 2013) as well as seed predators and fruit pulp consumers (Basset et al., 2021). Of the 36 species of herbivores and 2 species of seed predators found feeding on *P. limonensis* and the 21 species of herbivores, 1 species of seed predator, and 1 species of fruit pulp consumer found feeding on *P. marginata*, the 2 *Psychotria* species share 8 species of herbivores and none of the fruit antagonists in common (Sedio, 2013; Basset et al., 2021).

### Field collections

Sampling from 16 individual plants of each species, we collected expanding leaves, mature (fully-expanded) leaves, and ripe fruit from each plant in May-August 2019. All three sample types were collected from the same plant simultaneously. Expanding leaves were identified by their distal position on the branch and lighter color with respect to mature leaves and were collected when their leaf area reached approximately 50-70% of that of the neighboring mature leaves. Sufficient material was collected to yield at least 100 mg (dry mass) of each sample type, including 100 mg each of pulp (both species have soft exocarp which was combined with the mesocarp) and seeds, including the endosperm and embryo with lignified endocarp. All plants were located in qualitatively similar light environments in the forest understory, within 10 m of trails and with no canopy gaps larger than those resulting from trail maintenance.

Once collected, the tissues were immediately placed in a chilled and insulated cooler and taken back to the laboratory to be processed for extraction. Leaf samples were weighed and then placed in a −80°C freezer. Fruit samples were separated into pulp and seeds, weighed, and placed in −80°C freezer. With the low thickness of all sample types (intact fruit diameter ≤ 5 mm), freezing at −80°C was sufficiently rapid to prevent tissue damage due to ice crystal formation. After a minimum of 24 hours at −80°C, the samples were transferred to vacuum flasks and freeze-dried for 72 hours. After drying, samples were weighed and ground into powder, then returned to a −80°C freezer for storage until transport to Utah State University for secondary metabolite extractions.

### Secondary metabolite extractions and processing

#### Secondary metabolite extractions

Chemical extraction of plant secondary metabolites was carried out at Utah State University. For each sample, 80 mg of homogenized powder was weighed into a 1.8 mL polypropylene screw-top tube using a microbalance. To isolate the broadest possible range of phytochemicals, the extraction solvent used was 99.9% ethanol with 0.1% formic acid. Each sample was extracted with a total of 7.0 mL of the extraction solvent, conducted as five iterations of the following procedure. A glass syringe was used to add 1.4 mL of extraction solvent to the 1.8 mL tube containing the sample. The sample was mixed with the extraction solvent for 5 min in a vortexer at 1500 rpm and then centrifuged for 5 min at 12000 rpm, after which the supernatant was removed and added to a 20 mL glass scintillation vial. The supernatant from each of the five iterations was combined in the same 20 mL vial. The combined extract was dried at room temperature using a vacuum-centrifuge. The dried extract was then weighed and stored at −20° C until analysis.

#### UPLC-MS and post-processing

LC-MS data were collected using an Acquity I-class UPLC coupled to a Waters Synapt G2-S quadrupole time-of-flight mass spectrometer (Waters). For analysis, dried extracts were resuspended at 10 mg/mL in 75:25 water: acetonitrile + 0.1 % formic acid, with 2.0 μg/mL Stevioside as an internal standard. The extract was then sonicated for 10 min, after which a 20 μL aliquot was taken and diluted 10-fold with 75:25 water: acetonitrile + 0.1 % formic acid. The diluted aliquot was then vortexed and centrifuged (10 min,13,000 xg) and an aliquot (180 μL) was transferred to an LC-MS vial for analysis. Solvent blanks were injected at regular intervals during data collection. The autosampler temperature was 10°C and the injection volume was 1.5 μL. The column employed was a reverse-phase Acquity BEH C18 (2.1 mm ID x 150 mm, 1.7 um particle size, Waters) maintained at 35 °C at a flow rate of 0.2 mL/min. Solvent A was water with 0.1% formic acid and solvent B was acetonitrile with 0.1% formic acid (LCMS grade, Fisher Chemical). Solvent gradient: 0-0.5 min, 90% A; 0.5-1.0 min, 75% A; 1.0-8.0 min, 5% A; 8.0-10.0 min, held at 5% A; 10.0-11.0 min, 90% A; 11.0-15.0 min, held at 90% A.

Mass spectra and fragmentation spectra were collected separately in order to ensure accurate quantitation of molecular ion relative abundance. Fragmentation spectra were collected using data-dependent acquisition in positive-ion mode, with the following parameters: peak data recorded in centroid mode; 0.185 s MS scan time; 20-35 V collision energy ramp; argon collision gas; 125° C source temperature; 3 V capillary voltage; 30 V sample cone voltage; 350° C desolvation temperature; nitrogen desolvation at 500 L/hr; 10 μL/min lockspray flow rate; 0.1 s lockspray scan time; 20 s lockspray scan frequency; 3 lockspray scans to average; 0.5 Da lockspray mass window; 3 V lockspray capillary voltage. The lockspray solution was 1 ng/ μL leucine enkephalin, and sodium formate was used to calibrate the mass spectrometer.

Alignment, deconvolution, and annotation of molecular and adduct ions were conducted using methods described in Schneider et al. (2021).

#### Tandem mass spectrometry fragmentation data processing

The tandem mass spectrometry described above generated fragmentation spectra for all fragmentable molecular ions in our specialized metabolite extracts. These fragmentation spectra are diagnostic of molecular structure, and through pairwise comparison can be used to generate a network linking putative compounds to one another based on their structural similarity (Wang et al., 2016; Sedio et al., 2017; Aron et al., 2020). The methods used for this process are as described in Schneider et al. (2021) with the exception of an updated GNPS workflow: METABOLOMICS-SNETS-V2, version “release_30”.

## STATISTICAL ANALYSES

### Chemical richness across species and tissues

We compared the median and variance of chemical richness across the following groups: 1) among tissue types within each species, 2) among tissue types with species pooled within tissue type, and 3) between *P. limonensis* and *P. marginata* with tissue types pooled within each species. Unless otherwise stated, all tests were conducted using R statistical software (R Core Team, 2022). We first used a Shapiro-Wilk test to ascertain the normality of the richness data. We found the data to be left-skewed and non-transformable, and thus used the non-parametric Brown-Mood test to test for differences in median chemical richness among groups and post-hoc pairwise comparisons (controlling for the false discovery rate; Benjamini and Hochberg, 1995). These tests were conducted using the ‘coin’ (Zeileis et al., 2008) and ‘rcompanion’ (Mangiafico, 2022) R packages. To compare the variance of chemical richness, we used Levene’s test, found in the R package ‘car’ (Fox and Weisberg, 2019).

### Visualization and descriptive modeling of chemical compositional variance

We used partial least-squares discriminant analysis (PLS-DA) to reduce the dimensionality of the chemical compositional data in order to identify and quantify the principal components of variance with samples grouped by our categorical response variables of interest. Our groupings of interest were 1) tissue-grouped: tissue identity with species identity withheld from the model, 2) species-grouped: species identity with tissue identity withheld, and 3) tissue-with-species grouped models: both species and tissue identity assigned. This analysis was conducted through the MetaboAnalyst version 5.0 online workflow (Xia and Wishart, 2016).

### Chemical compositional similarity across species and tissues

Metabolome-scale chemical similarity among samples and groups of samples was quantified using two complementary distance-based metrics. First, we calculated pairwise Bray-Curtis chemical compositional similarities (BCS) of all samples using the R statistics package ‘vegan’ (Oksanen et al., 2022). In this context, the Bray-Curtis index quantifies similarity based on the number of compounds that are shared between samples and the relative ion abundances of each shared compound (Sedio et al., 2017; Schneider et al., 2021). Second, we used the GNPS molecular networking output described above and an R-based workflow adapted from Sedio et al. (2018) to calculate the pairwise molecular structural similarities of all samples, referred to as chemical structural similarity (CSS). This workflow is described in detail in Schneider, et al. (2021). Briefly, for each sample-pair comparison, the samples’ structural similarity is calculated by taking the mean of all pairwise compound-level molecular fragment similarity comparisons. These nested pairwise comparisons are used to generate a sample-by-sample distance matrix similar in format to the Bray-Curtis index.

We compared the means of BCS and CSS across groups at four distinct levels dictated by potential sources of biological variation: intra-specific intra-tissue, intra-specific inter-tissue, inter-specific intra-tissue, and inter-specific inter-tissue. To test for differences in group means, we used one-way permutational ANOVA tests in the R package ‘lmPerm’ (Wheeler and Torchiano, 2016), with Tukey’s HSD for post-hoc pairwise tests.

We then tested the effects of species, tissue type, and individual plant identity on the distribution of BCS using a series of PERMANOVA model tests (Oksanen et al., 2022). First, we tested the effects of species, tissue, and their interaction on chemical compositional similarity using a fixed-effects model. Next, we tested whether tissue-level variation was nested within plant-level variation for each species by comparing fixed-effect models—with tissue as the only fixed effect—to mixed-effect models with plant ID added as a random effect (separate models for *P. limonensis* and *P. marginata*). Finally, we tested the effect of plant-level variation on chemical compositional similarity within each species, with plant ID as a fixed effect and species as a random effect in a mixed-effect model. In all models, the number of permutations used was n = 999. Individual plants for which any of the four tissues were not collected (due to lack of vegetative or reproductive growth) were omitted from the latter two analyses. PERMANOVA assumes that the samples are exchangeable under a true null hypothesis and is sensitive to differences in multivariate dispersion (Anderson, 2001, 2017). We used a permutation test (N = 999 permutations, function ‘betadisper’ in the R package ‘vegan’) to quantify homogeneity of multivariate dispersion from the overall centroid, an analogue of Levene’s tests for homogeneity of variances, across the four tissue types followed by a post-hoc Tukey HSD test to assess pairwise differences in multivariate location and dispersion between tissue types. All PERMANOVAs were conducted using the function ‘adonis2’ in the R package ‘vegan’ (Oksanen et al., 2022).

CSS data were omitted from PERMANOVA tests, as these data were characterized by presence/absence of compounds. Given the low levels of variation in compound presence/absence established in our preliminary analyses, CSS data were not informative of intra-group variation. In addition, the homogeneity of multivariate dispersion test was not conducted for CSS due to the reduced sample size from which molecular structural data were collected.

## RESULTS

### Chemical richness across species and tissues

Across all sample groups—expanding leaves, mature leaves, ripe pulp, and ripe seeds from *Psychotria limonensis* and *Psychotria marginata*—our UPLC-MS analysis yielded 2722 molecular features annotated as putative compounds, hereafter referred to as compounds. Combining the two species to summarize tissue-level results, we found that 48.6% of the 2722 compounds were present in all four tissue types, while 18.7% occurred only in expanding and mature leaves, 8.4% occurred only in fruit pulp and seed, and the remaining 24.3% occurred in a subset of both fruit and leaf tissues (**Fig. 1**).

**Figure 1.**
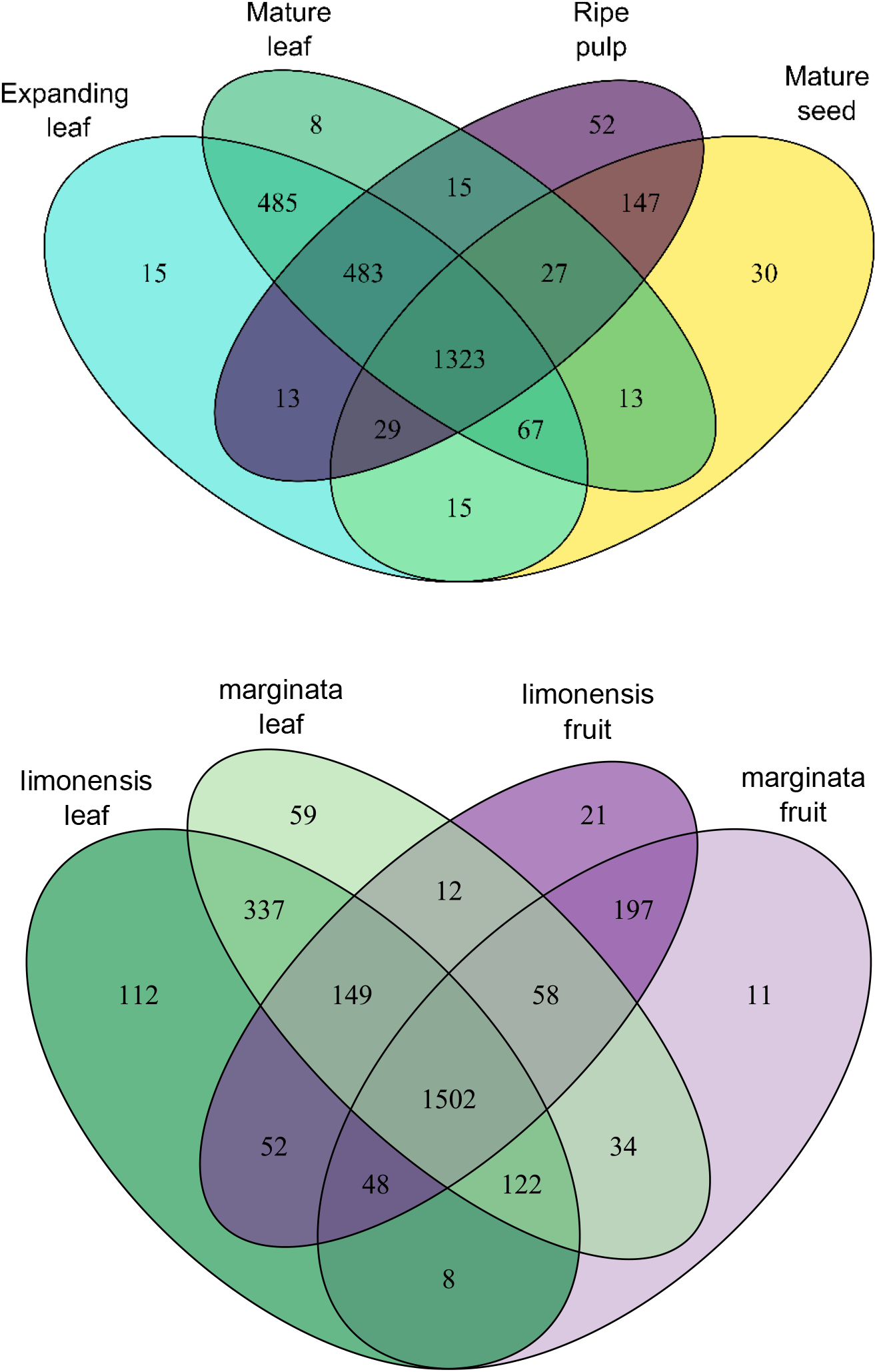
Venn diagrams of specialized compound richness. Top: compound richness by tissue identity with species combined. Bottom: compound richness by species and organ identity, with expanding and mature leaves combined into “leaf” and pulp and seed combined into “fruit”.

We found no significant difference in median chemical richness between the two species overall (pooling tissue types within species; Z = −0.0928, P = 0.926; **Fig. 2**). However, comparing among tissue types with species pooled, we found a significant difference in tissue-level medians overall (χ^2^ = 93.93, df = 3, P < 0.001) and, in each pairwise comparison, except for that of the two leaf development stages (expanding leaf – mature leaf: P = 0.695; all other comparisons: P < 0.001; **Fig. 2**).

**Figure 2.**
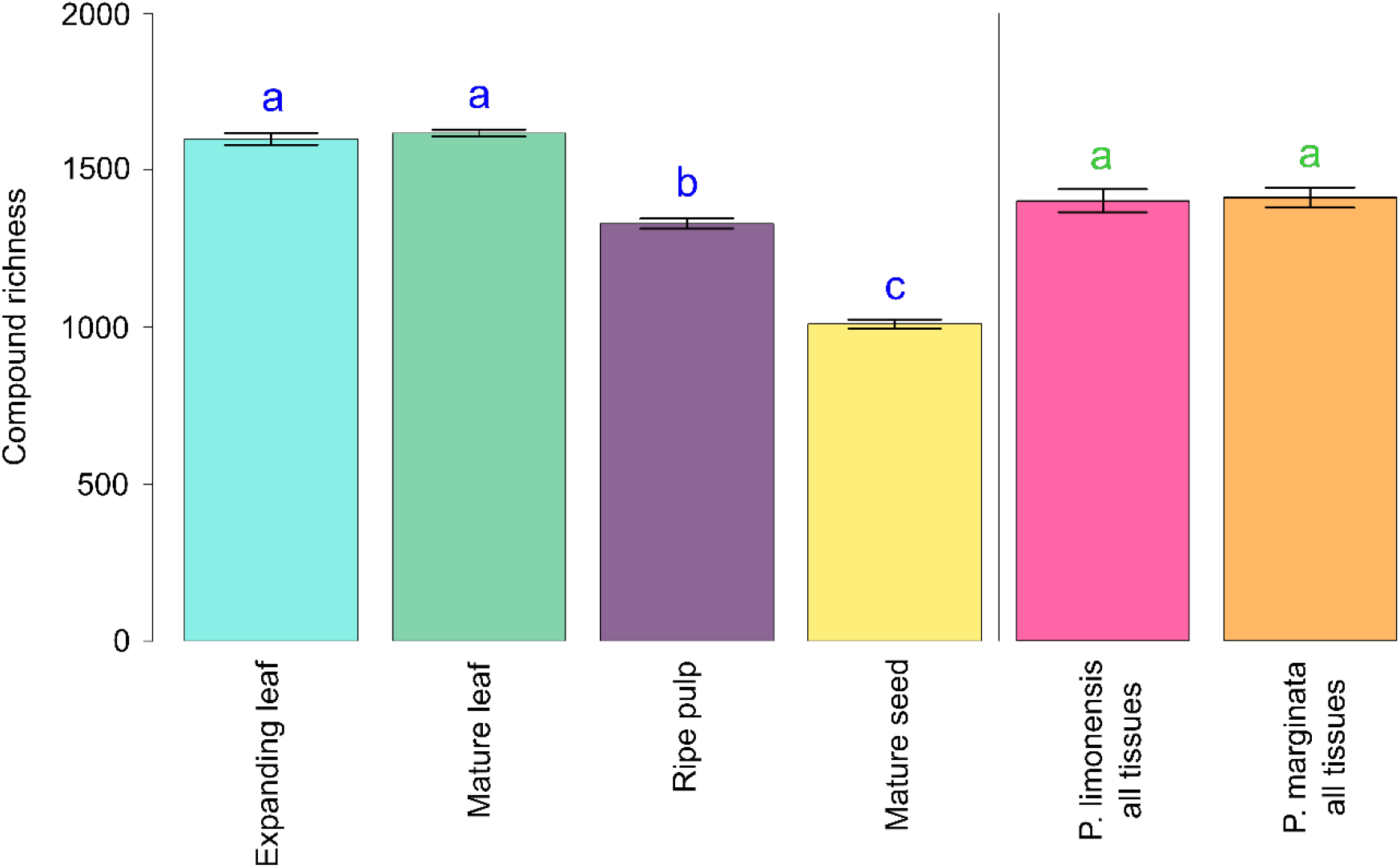
Group-level variation in compound richness. Left: Compound richness by tissue types with species pooled within tissue type; right: Compound richness by species with tissues pooled within species. Significant differences (P ≤ 0.05) between groups are indicated by non-shared letters above error bars. Error bars denote mean +/− SEM.

Using Levene’s test to evaluate the homogeneity of variance in compound occurrence across all samples within each tissue, we found somewhat significant heterogeneity of variance among the four tissues overall (F_3,109_ = 2.82, P = 0.042); however, we found no significant heterogeneity in any post-hoc pairwise comparison of tissues (**Table 1**). At the species level, a Levene’s test indicated that variance in compound occurrence between *P. limonensis* and *P. marginata* was marginally heterogeneous (F_1,111_ = 3.71, P = 0.057).

**Table 1.** Results of Levene’s tests for homogeneity of variance across groups conducted with each pairwise combination of tissues.

|  | Expanding leaf | Mature leaf | Ripe pulp |
| --- | --- | --- | --- |
| Expanding leaf | -- | $F_{1,26} = 2.44$<br>$P = 0.131$ | $F_{1,24} = 1.12$<br>$P = 0.300$ |
| Mature leaf | $F_{1,26} = 2.44$<br>$P = 0.131$ | -- | $F_{1,29} = 0.006$<br>$P = 0.942$ |
| Ripe pulp | $F_{1,24} = 1.12$<br>$P = 0.300$ | $F_{1,29} = 0.006$<br>$P = 0.942$ | -- |
| Mature seed | $F_{1,24} = 1.27$<br>$P = 0.271$ | $F_{1,29} = 0.003$<br>$P = 0.955$ | $F_{1,24} = 1.1837$<br>0.2874 |

### Visualization and descriptive modeling of chemical compositional variance

Our next set of analyses examined the patterns of variance in the chemical compositional data as well as the factors influencing these patterns. These data comprise the sample-level occurrence and relative ion abundance of the 2722 compounds described above, with samples grouped by tissue and/or species depending on the analysis.

We constructed three PLS-DA models for each of the different groupings: species-grouped model, tissue-grouped model, and tissue-with-species grouped models (**Fig. 3**). In the species-grouped model, which maximized chemical compositional variance between species, the primary component “component 1” accounted for only 11.9% of total variance and did not produce a statistically significant separation distance between the two groups (Bohn-Wolfe permutation test, p = 0.176, nperm = 1000). Further, the first component in this model explains 7.3% less total variance in the data than does the second component, which explained 19.2% of the total variance. The PLS-DA model by definition sets component 2 to be orthogonal to component 1. As observed in **Fig. 3**, component 1 reflects interspecific variance in chemical composition, whereas component 2 reflects intraspecific variance in chemical composition. Thus, the species-grouped model suggests that intraspecific variance is greater than interspecific variance in chemical composition across *P. limonensis* and *P. marginata*. In contrast to the species-grouped model, the tissue-grouped and tissue-with-species grouped models both yielded first components accounting for more total variance than each model’s subsequent components (**Fig. 3**). In addition, the tissue-with-species grouped model produced a first component with loadings that showed greater separation between vegetative and reproductive tissue types than between species (**Fig. 3**). Finally, both of the latter models produced a statistically significant separation distance between groups (both models: p < 0.001, nperm = 1000).

**Figure 3.**
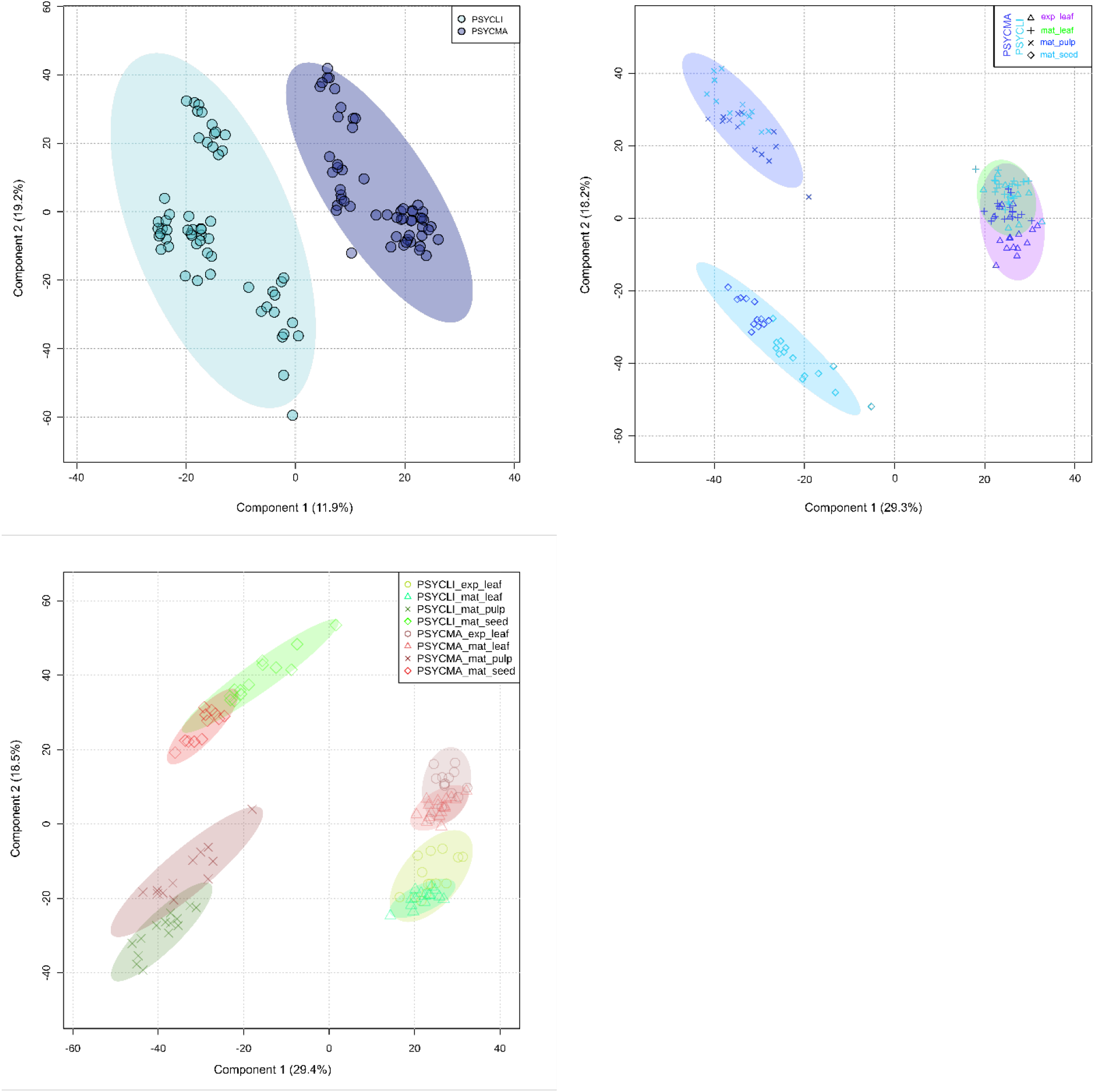
PLS-DA principal component plots for chemical composition by species (top left), tissue (top right), and tissue-with-species (bottom left). Shading indicates 95% confidence intervals of group variance.

### Chemical compositional similarity across species and tissues

Overall, the permutational ANOVA indicated highly significant differences in mean similarities among group levels (P < 0.001). In pairwise tests, comparing the pairwise Bray-Curtis chemical compositional similarity (BCS) of each tissue within and across the two *Psychotria* species (intra-specific inter-tissue and inter-specific inter-tissue), we found that all four tissues had significantly higher mean similarity within species than between species (P < 0.001) (**Fig. 4**). In intra-specific inter-tissue comparisons, the two vegetative tissues were more similar to each other within species than to either of the reproductive tissues within each species (P < 0.001), as were the two reproductive tissues with respect to each other and the vegetative tissues (P < 0.001). In inter-specific intra-tissue comparisons, the two vegetative tissue types exhibited lower mean interspecific similarity between species than did the two reproductive tissue types between species (P < 0.001) (**Fig. 4**).

**Figure 4.**
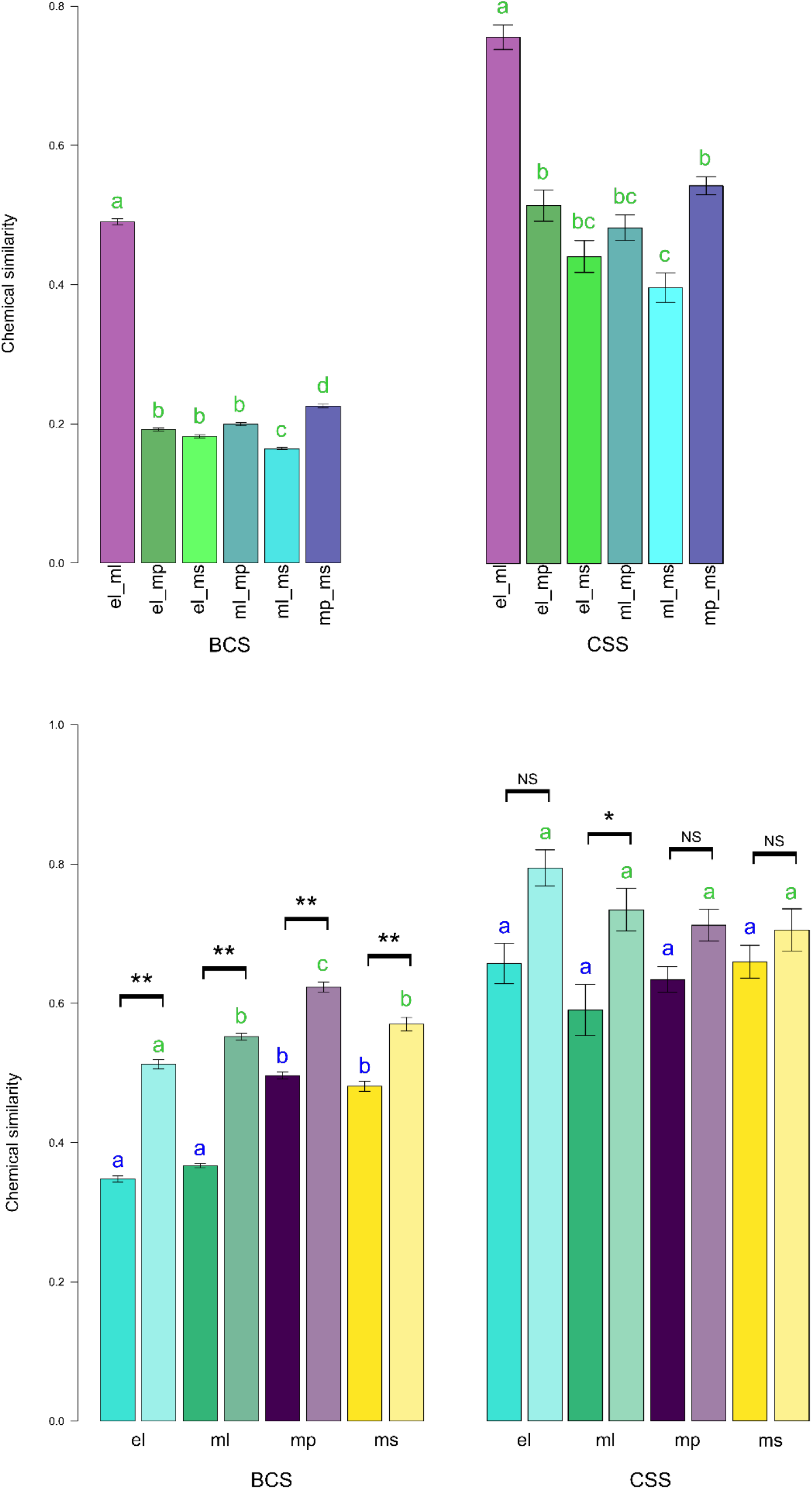
Bray-Curtis (BCS) and chemical structural (CSS) similarity of samples within and across species and tissues. Tissues are coded as follows: el = expanding leaf, ml = mature leaf, mp = mature pulp, ms = mature seed. Error bars denote mean +/− SEM. **Top**: intra-species inter-tissue comparisons; significant differences (P ≤ 0.05) between groups are indicated by non-shared letters above error bars. **Bottom**: intra-tissue comparisons; inter-species in dark shading and intra-species in light shading. Significant differences (P ≤ 0.05) between groups are indicated by non-shared letters above error bars, with blue letters for comparisons among inter-species groups and green letters for comparisons among intra-species groups. Brackets above error bars indicate statistical significance of inter-species vs. intra-species comparison for each tissue type; NS = not significant, * = P ≤ 0.05, ** = P < 0.001.

In comparisons paralleling those described for BCS, chemical compositional similarity (CSS) exhibited trends largely similar to BCS. These analogous trends were the case overall, with strong statistical support from the permutational ANOVA (P < 0.001), as well as in pairwise tests, though statistical support was weaker in some cases (**Fig. 4**). The main exception was with respect to differences in intra-tissue similarity at inter- and intra-specific group levels, which in CSS were equivalent among vegetative and reproductive tissue types. (**Fig. 4**).

Testing the effects of species, tissue, and their interaction on BCS using PERMANOVA demonstrated that both factors, along with their interaction, had significant effects on chemical compositional similarity. Tissue type (F_3,112_ = 47, R^2^ = 0.49, P < 0.001) explained six-fold or more of the variation in chemical compositional similarity than did species (F_1,112_ = 22, R^2^ = 0.07, P < 0.001) or the tissue-by-species interaction (F_3,112_ = 7.9, R^2^ = 0.08, P < 0.001). Examining the nesting of tissue-level within plant-level variation, we found that nested models were not discernible from non-nested models in either species (*P. limonensis:* F_3,31_ = 18, R^2^ = 0.66, P < 0.001; *P. marginata:* F_3,35_ = 19, R^2^ = 0.65, P < 0.001), indicating a negligible level of nesting for tissue-level BCS variation within plant-level BCS variation. Indeed, when examined as a fixed effect, plant-level variation itself did not contribute significantly to BCS trends within species (F_16,67_ = 19, R^2^ = 0.17, P > 0.999).

Following up our PERMANOVA tests with analyses of BCS variance across tissues with species pooled within tissue type, we found significant heterogeneity of the corresponding multivariate dispersions (F_3,109_ = 9.95, P < 0.001). Pairwise Tukey’s HSD tests indicated that both expanding and mature leaves exhibited significantly more chemical compositional dispersion than did ripe pulp or mature seeds (**Fig. 5**). Within leaves, however, expanding and mature tissues did not differ significantly from one another in chemical compositional dispersion (P = 0.95). Similarly, within fruit, the dispersions of BCS among ripe pulp and mature seeds were comparable (P = 0.78).

**Figure 5.**
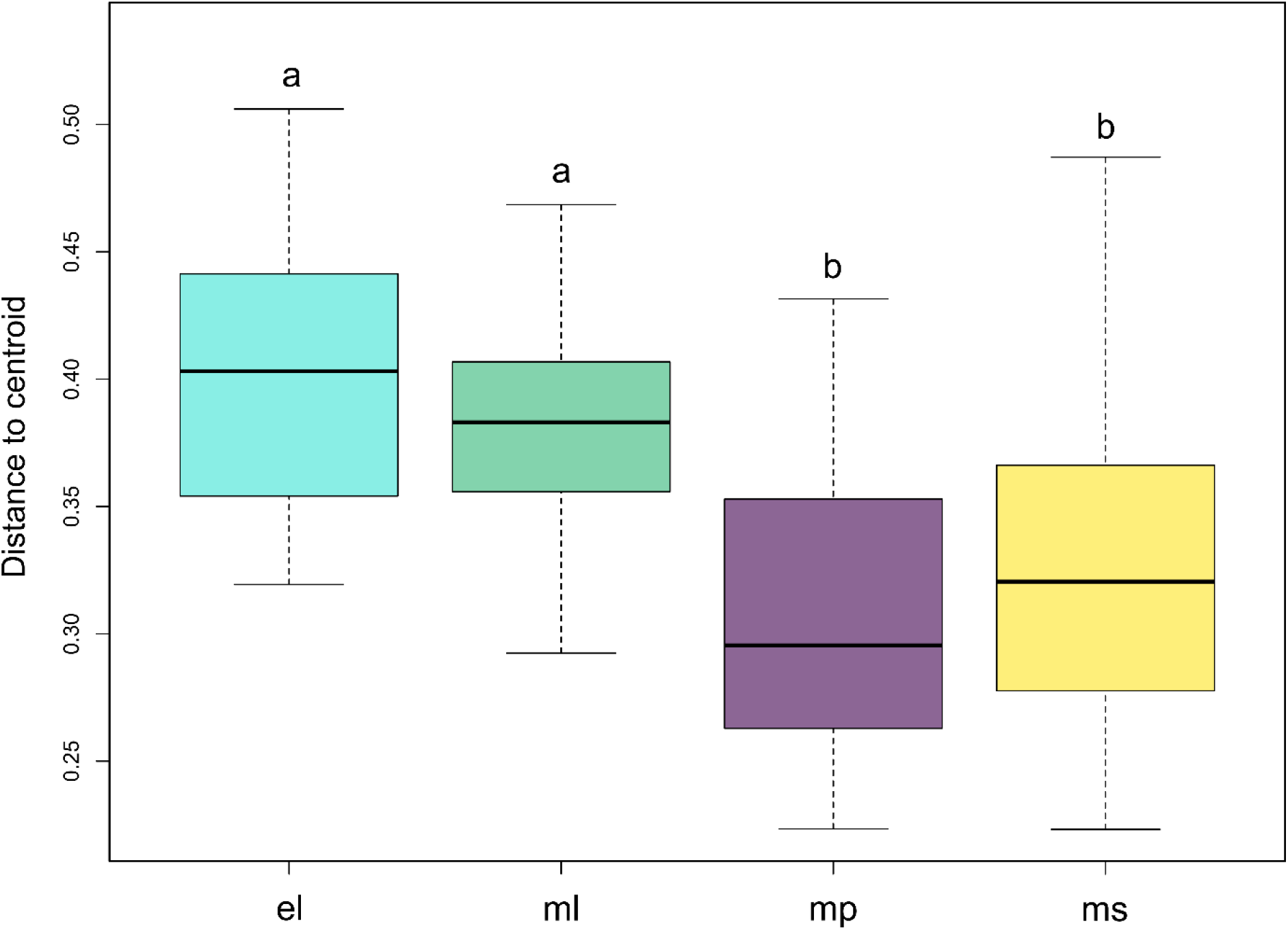
Comparison of heterogeneity of chemical composition by tissue. Species are pooled within tissue type. Tissue coding is as in Figure 4. Significant differences (P ≤ 0.05) between groups are indicated by non-shared letters above error bars. Boxes indicate median +/− IQR. Whiskers indicate spread of data within 1.5x IQR.

## DISCUSSION

Zoochorous fruit and their inherent diversity of biotic interactions have gained increasing attention from researchers seeking an organismal perspective on the ecology and evolution of plant specialized metabolites (Schneider et al., 2021; Whitehead et al., 2021). Numerous case studies comparing organs of a given species have revealed or corroborated facets of organ-specific specialized metabolite composition (reviewed in Kessler and Halitschke, 2009; Whitehead et al., 2013; Berardi et al., 2016; Schneider et al., 2021). These studies, as well as cases of pleiotropy where key metabolites are shared among organs (Adler et al., 2012; Keith and Mitchell-Olds, 2019), have evidenced organ-level selective pressures as a key evolutionary dynamic in yielding the overall species- and community-level diversity of plant specialized metabolites. Our case study builds on those preceding, adding to the evidence of inter-organ differences in chemical richness and variance as well as the presence of organ-specific metabolites. Perhaps most notably, our study is the first to document the scale of inter-organ variation in specialized metabolites in comparison to inter-individual variation within species, with the former greatly exceeding the latter. Finally, our study confirms our system-specific hypotheses of specialized metabolite composition based on the Interaction Diversity Hypothesis.

Using untargeted metabolomics, we examined the biological patterns of variation exhibited by specialized metabolites across leaves and fruit of two Neotropical *Psychotria* species. As a whole, our comparison of leaf- and fruit-associated specialized metabolites indicates that chemical variation between vegetative and reproductive organs, both within and between species, dwarfs analogous variation across all other biological scales represented in our study. This was the case for chemical variation as quantified by richness of compounds, PLS-DA multivariate main components, compositional similarity, and chemical structural similarity. Such a pattern of variation is consistent with the specialized metabolite composition of each organ being shaped – at least in part – by the biotic interactions specific to that organ.

If we are to regard the specialized metabolite composition of each organ as a distinct multivariate trait, it must first be established whether these putative traits 1) occupy distinct neighborhoods of trait space and 2) vary independently of one another across individuals. With a sufficient sample size and high-resolution chromatography and mass spectrometry, we were able to compare metabolome-scale trait values and intra- and inter-organ variation of specialized metabolite composition. We quantified multivariate specialized metabolite composition using the complementary methods of PLS-DA dimension reduction (**Fig. 3**) and distance based BCS and CSS (**Fig. 4**). PLS-DA confirmed that the specialized metabolite composition of leaves, fruit pulp, and seeds all occupy different neighborhoods of multivariate trait space (**Fig. 3**). BCS and CSS confirmed that specialized metabolite composition was less similar among different organs of a given species than across replicates of a given organ across individuals and even across species (**Fig. 4**). Finally, applying PERMANOVA testing to the BCS data, we were able to confirm that plant identity had no significant influence on organ-level variation in chemical composition, indicating that leaf and fruit chemical composition varied independently of one another regardless of whether the samples were taken from the same plant.

Regarding our system-specific, IDH-based hypotheses, the chemical compositional characteristics of leaves and fruit were sufficiently distinctive to allow informative contrasts. Taken together, our quantitative measures of specialized metabolite relative abundances and molecular structural relationships showed that leaves had a larger number of unique chemical compounds as compared to fruit (**Fig.1**) as well as higher degrees of variation in metabolite relative abundances among individuals and between species (**Figs. 4 and 5**). Both of these results supported our predictions, attributable to the higher richness and taxonomic diversity of interactions in which leaves are involved as compared to fruits in these *Psychotria* as well as the predicted costs of chemical variation to consumers.

While fruit tissues had fewer unique chemical compounds than leaves, the number of compounds found only in fruits was still in the hundreds (**Fig.1**). Further, the interspecific chemical structural variation was comparable to that of leaves (**Fig. 4**), indicating that leaf-specific and fruit-specific metabolites have similar degrees of structural divergence with respect to ubiquitous metabolites. Despite the lower taxonomic richness and phylogenetic diversity of fruit-centric interactions as compared leaf-centric interactions in this system, these results are consistent with IDH. We interpret interaction diversity as reflecting the strength and sign of interactions as well as the richness and diversity of the species involved. While quantifying the strength of the antagonistic and mutualistic interactions in this system is beyond the scope of our study, the fact that fruit are involved in both antagonistic and mutualistic interactions suggests that fruit are exposed to substantial interaction diversity as reflected in the selective pressures on fruits’ specialized metabolites. Developing the mathematical underpinnings of IDH to quantitatively incorporate the aspects of interaction diversity which we have discussed will represent an important component of future work in this field.

## CONCLUSIONS

A growing body of evidence, to which we contribute with the present study, demonstrates ever more clearly that zoochorous fruit and leaves can represent distinct compartments in the ecology and evolution of plant specialized metabolites. Such partitioning of chemical composition based on localized selective pressures is likely applicable to other ecologically distinct organs and tissues as well, such as flowers (Berardi et al., 2016) and roots (Wang et al., 2015), and our ongoing research seeks to integrate all such compartments into a whole-plant perspective. The genetic and phylogenetic dynamics of the genomic connections of these compartments remain unelucidated, but recognizing specialized metabolite composition as a set of associated organ- or tissue-level traits is a crucial step toward understanding their micro- and macroevolutionary basis.

## ACKNOWLEDGEMENTS

This research was supported by the National Science Foundation grant no. IOS-1953934 to NGB; start-up funds from Utah State University to NGB; the Richard J. and Marion A. Shaw Scholarship and the Gene and Ruth Miller Scholarship from the Department of Biology at USU to CAC, University Honors Program funds at USU to CAC, and an Undergraduate Research and Creative Opportunity Grant from the Office of Research at USU to CAC.

## AUTHOR CONTRIBUTIONS

CAC, NGB, and EMJ designed the field study. CAC collected field samples and conducted chemical extractions. GFS designed and conducted chemical analyses. GFS designed and conducted the statistical analyses with contributions from NGB. GFS led the writing of the initial draft with contributions from NGB. All authors contributed to revising the manuscript.

## DATA AVAILABILITY STATEMENT

UPLC-MS molecular network data will be publicly available through GNPS (gnps.ucsd.edu). All other data and R code will be publicly available through GitHub. Hyperlinks to repositories will be added upon manuscript acceptance after peer review. Data will be available upon request during peer review.

